# ArtSeg: Rapid Artifact Segmentation and Removal in Brightfield Cell Microscopy Images

**DOI:** 10.1101/2022.01.24.477467

**Authors:** Mohammed A. S. Ali, Kaspar Hollo, Tõnis Laasfeld, Jane Torp, Maris-Johanna Tahk, Ago Rinken, Kaupo Palo, Leopold Parts, Dmytro Fishman

## Abstract

Brightfield cell microscopy is a foundational tool in life sciences. The acquired images are prone to contain visual artifacts that hinder downstream analysis, and automatically removing them is therefore of great practical interest. Deep convolutional neural networks are state-of-the-art for image segmentation, but require pixel-level annotations, which are time-consuming to produce. Here, we propose ScoreCAM-U-Net, a pipeline to segment artifactual regions in brightfield images with limited user input. The model is trained using only image-level labels, so the process is faster by orders of magnitude compared to pixel-level annotation, but without substantially sacrificing the segmentation performance. We confirm that artifacts indeed exist with different shapes and sizes in three different brightfield microscopy image datasets, and distort downstream analyses such as nuclei segmentation, morphometry and fluorescence intensity quantification. We then demonstrate that our automated artifact removal ameliorates this problem. Such rapid cleaning of acquired images using the power of deep learning models is likely to become a standard step for all large scale microscopy experiments.

## 1. Introduction

Advanced microscopes extract rich visual information from biological samples at scales from individual atoms to cells and tissues. Among the different imaging modalities, brightfield illumination with transmitted light is the simplest to acquire while avoiding damaging the sample^1^. The usefulness of this technology has led to its widespread adoption^2–4^, and thereby to a dramatic increase in the volumes of microscopy data. However, the automated analysis techniques required to extract information at scale are often hindered by the artifacts present in the images^5,6^. Detecting and neutralizing the impact of such problematic image areas would provide more accurate results from experiments^3^, making artifact segmentation an important, albeit overlooked, research area in cell biology and beyond^7,8^.

While any signal that deviates from the reflection of expectation can be considered artifactual^9^, the common source of artifacts in cell microscopy is the introduction of foreign objects during sample preparation. These include dust, fragments of dead cells, bacterial contamination, reagent impurities, defects on the light path, etc. We focus on detecting these low-level anomalies^8,10^ in brightfield microscopy and use the term *artifact* with this meaning. Manually identifying all the affected images or image regions is a time-consuming solution to this problem^11,12^. A common alternative approach for large datasets is computer-aided delineation and removal of the artifacts, but two complexities make this task challenging. First, artifacts are generated stochastically leading to sparse data. Second, artifact characteristics, such as morphology and texture, are often very heterogeneous and rarely well defined. These features render computational modeling difficult.

Deep learning has emerged as the favored solution to artifact detection^7,8^. While strongly supervised convolutional neural networks (CNN) such as U-Net^13–17^ are state-of-the-art for most computer vision tasks, they cannot overcome some challenges that artifact detection brings^7^. A major bottleneck for the strongly supervised deep learning methods is their requirement of pixel-level annotation, which is time-consuming, and requires substantial expertise. As an alternative, weakly supervised techniques such as ScoreCAM^18^, which involve only image-level labeling, greatly reduce the time needed to prepare the dataset. In particular, generative autoencoder-based models^19–24^ are trained to reconstruct artifact-free images and report artifacts on test images as areas with large reconstruction error. Alternatively, one-class classification approaches^25,26,27^ train a classifier on artifact-free images and report artifacts as images with a low probability of belonging to this clean class. However, neither method performs well enough for adaptation in routine microscopy image processing workflows. Combining the performance advantages of the strongly supervised methods and the convenience of image-level annotations would therefore be of great practical interest and impact.

Here, we combine the merits of weakly and strongly supervised methods for artifact segmentation from brightfield cell microscopy images using only image-level annotations. To our knowledge, this is the first attempt to segment artifacts in microscopy images in a weakly supervised way. We introduce ScoreCAM-U-Net, a model that combines the informative pixel-level^4^ and cheap-to-generate image-level^18^ annotation schemes, and accurately detects artifacts in held-out samples. As training is performed using only image-level labels, generating training data is orders of magnitude cheaper, but without substantially sacrificing performance compared to pixel-level data. We confirm that artifacts in microscopy images confound downstream analyses such as nuclei segmentation or quantification of ligand binding, and demonstrate that ScoreCAM-U-Net successfully overcomes these problems.

## 2. Methods

To delineate artifacts from brightfield microscopy images, we introduce ScoreCAM-U-Net, a method that uses image-level annotations as input for training, and produces artifact segmentations as an output. We compare the performance of our pipeline with a strongly supervised counterpart trained on pixel-level annotations as well as with state-of-art models that are trained using image-level labeling on three different datasets.

### 2.1. Datasets

We chose three datasets for this study to cover multiple common variables in experimental design to better assess the generalizability of the results. Overall, the datasets cover nine different cell lines, fixed and live cells, two different plate formats and two microscopes. The datasets provenances have been described previously^3,28,4,29^ and we briefly describe their most important properties here.

#### Seven cell lines dataset

Seven types of cells including human cells from breast cancer (MCF7), fibrosarcoma (HT1080), cervical cancer (HeLa), hepatocellular carcinoma (HepG2), alveolar basal epithelial (A549), dog cells from kidney tissue (MDCK), and mouse embryonic fibroblast cells (NIH3T3) were seeded in Collagen type 1-coated CellCarrier-384 Ultra Microplates (PerkinElmer, Waltham, MA; cat. 6057700). The cells were stained with 10μg/ml Hoechst 33342 (Thermo Fisher, Waltham, MA; cat. H3570) and fixed in formaldehyde (Sigma, St. Louis, MO; cat. 252549). A 20x water immersion objective was used to acquire images on an Opera Phenix™ high-content screening system (PerkinElmer) in confocal mode. Nine fields of view were acquired from each well with a total of 3024 images of size 1080×1080px (1px = 0.59μm) with 350 cells in each field of view on average. All fields of view were imaged in fluorescent and brightfield modalities, with one modality acquired first on all wells and then the second. This dataset is referred to as “seven cell lines” in the further text.

#### LNCaP dataset

The cells of human prostate adenocarcinoma (LNCaP, from ATCC) were seeded in a CellCarrier-384 Ultra Microplate (PerkinElmer), fixed in formaldehyde, and stained using DRAQ5 fluor (Abcam, Cambridge, United Kingdom) to tag nuclear DNA. A 20x objective was used to acquire images on a CellVoyager 7000 (Yokogawa, Tokyo, Japan) instrument in confocal mode to acquire fluorescence and brightfield images of size 2556 × 2156 pixels (1 pixel = 0.325 μm) with 681 cell in each field of view on average. Similar to the seven cell lines dataset, one modality was acquired on all wells before moving on to the second modality.

#### ArtSeg-CHO-M4R dataset

The imaging was performed as described previously^28^. Briefly, live CHO-K1-hM_4_R cells were seeded with a density of 25 000 cells per well into μ-Plate 96 Well Black plate (Ibidi) 5-7 hours before the imaging to allow attachment. All the experiments were performed in the cell culture medium DMEM/F-12 with 9% FBS (Sigma), antibiotic antimycotic solution (100 U/ml penicillin, 0.1 mg/ml streptomycin, 0.25 μg/ml amphotericin B, Sigma) and 750 μg/ml of selection antibiotic geneticin (G418, Capricorn Scientific). The final volume in the well was 200 μl. All imaging experiments were carried out at 37 °C in the 5% CO_2_ atmosphere. The images were captured with Cytation 5 Imaging Multi-Mode Reader (BioTek, Bad Friedrichshall, Germany). Images were obtained using a LUCPLFLN 20x objective lens with working-distance of 6.6 mm, and numerical aperture of 0.45 (Olympus), using LED excitation source with 351(40) nm filter and captured with 593(40) nm emission filter. The field of view size was 1224 x 904 pixels (1 pixel = 0.323 μm). For a single field of view, a brightfield image was obtained first, which was immediately followed by fluorescence image acquisition. These steps were repeated for four fields of view in each well. In all experiments, a constant concentration of 2 nM UR-CG072^30^, a TAMRA labeled fluorescence ligand was used to visualize cells expressing muscarinic M4 receptors in the fluorescence channel. In concentration-response experiments atropine, arecholine (Sigma), UNSW-MK259^31^ and UR-SK75^32^ were used. UNSW-MK259, UR-SK75 and UR-CG072 were kindly provided by Dr. Max Keller from the University of Regensburg. The ArtSeg-CHO-M4R dataset is made freely available for public use.

#### Artifact annotation

The seven cell lines and LNCaP data were inspected and 11.4% and 6.5% of the samples were found to have artifacts, 344/3024 and 51/784 fields-of-view respectively. The same number of fields-of-view from each dataset were randomly sampled to be used as training images without artifacts. At the same time, 99.2% of samples in the ArtSeg-CHO-M4R dataset (1171/1181) were found to have artifacts. The clean images for this dataset were generated as described below.

For all three datasets, pixel-level ground truth masks of artifacts were generated by manual annotation. All annotators had prior training in bioimage analysis, microscopy and cell biology. For seven cell lines and LNCaP datasets, the artifacts were annotated as polygons using VGG image annotator^33^ and for ArtSeg-CHO-M4R dataset, as freehand annotations with the MembraneTools module of Aparecium software^34^. For all datasets, the artifact pixels were annotated while keeping the number of background pixels annotated as artifacts as low as possible. For the ArtSeg-CHO-M4R dataset, the artifact annotations contain a considerable number of background pixels in some images as it speeds up annotation and better reflects the annotation process in real-world conditions.

For obtaining the weak labels for the seven cell lines and the LNCaP datasets, the images were classified into either clean or artifact-containing images after a brief inspection by the annotator. For the ArtSeg-CHO-M4R dataset, as the vast majority of images contain at least one artifact, the clean images were generated by replacing the pixel values of manually annotated artifacts with the values of the corresponding pixels in the estimated background image. The background is estimated by fitting the original image with a two-dimensional second order polynomial function^35^. To simulate imaging noise, a zero-centered noise profile of the background pixels is added to the estimated background. No artifacts could be detected from the resulting images. The modified areas were also not visually detectable by human experts.

### 2.2. ScoreCAM-U-Net for artifact segmentation

Our weakly supervised artifact segmentation pipeline combines the ScoreCAM model^18^ that highlights areas of the image most useful for differentiating between clean and artifact-containing images with U-Net model^4^ that directly classifies pixels into categories. We call this pipeline “ScoreCAM-U-Net” (Figure 1).

**Figure 1:**
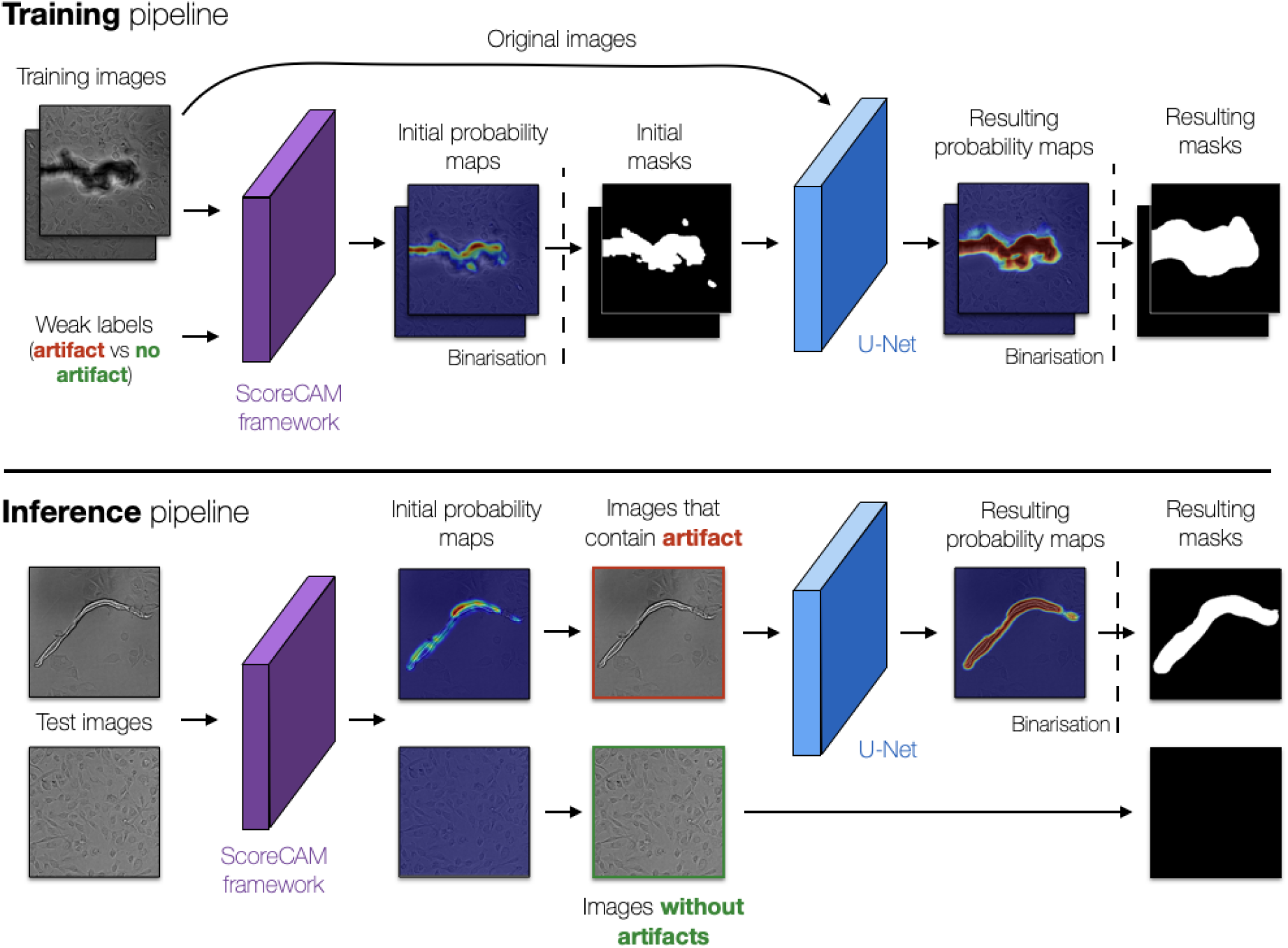
Artifact segmentation pipeline - ScoreCAM-U-Net. During training (top), ScoreCAM^18^ (purple) is used to generate pixel-level probability maps of artifacts and the corresponding binary masks that are used to train the U-Net^4^ segmentation model (blue). During the inference (bottom), the trained U-Net (blue) is used to segment artifacts from the images that were deemed to contain artifacts (image with red borders) by the ScoreCAM (purple). Vertical dashed lines: binarization of pixel probability maps values.

ScoreCAM^18^ is a technique used to explain predictions made by deep learning methods, mostly applied to models that perform image classification. ScoreCAM analyzes both the model output and the corresponding image, and highlights parts of the image that had a large impact for the particular prediction. It proceeds in four steps. First, visual representations (activation maps) of the last convolutional layer are extracted from an image classification model (ResNet^36^ in our implementation). Next, each activation map is upscaled to match the size of the input image, normalized to a range between 0 and 1, and projected onto a copy of the input image via multiplication, producing a projected input image. Then, the classification model (ResNet in our implementation) uses projected inputs to calculate the probability of the input image belonging to each class. Finally, all activation maps are summed, each multiplied by the corresponding class-largest probability and passed through the ReLU^37^ activation function to generate the final output (Supplementary Figure 1). Unlike other competitors that rely on gradients, ScoreCAM uses the largest class probability to obtain the resulting map. It has been empirically shown that this feature makes ScoreCAM less noisy and therefore more useful in practice^18^.

The strongly supervised U-Net^13^ model has already been successfully adapted for brightfield nuclei segmentation and its architecture is described in detail in the corresponding paper^4^ (Supplementary Figure 1). The architecture consists of an encoder and a decoder connected by a bottleneck, and skip links which pass the signal from the encoder to the decoder. We used an encoder consisting of 15 convolutional layers that use convolutional filters of size 3×3 and a rectified linear unit (ReLU)^4,37^ activation function. After every third layer, there is a 2×2 max-pooling layer and a skip connection to the decoder. Symmetrically, the decoder has 15 convolutional layers with ReLU activation functions. After every third convolutional layer, there is an upsampling layer that upscales its input height and width by a factor of 2. Finally, the bottleneck after the encoder has three convolutional layers. There are 64 filters in each convolutional layer in the encoder, decoder, and bottleneck.

### 2.3. Model training and evaluation

#### 2.3.1. Training

To train the ScoreCAM-U-Net model, the ResNet50^36^ classification model in the ScoreCAM^18^ framework was first trained to classify clean and artifact-containing images. The model is trained on the seven cell lines dataset using 482 images for training, 101 for validation and 104 for testing; on ArtSeg-CHO-M4R, using 1386 images for training, 404 for validation, and 572 for testing; and on LNCaP, using 70 images for training, 16 images for validation and 16 images for testing. The test set in ArtSeg-CHO-M4R dataset was chosen such that ten concentration-response curves with multiple competitive ligands could be obtained. The Adam optimizer^38^ was used to optimize binary cross-entropy loss for 150 epochs. The initial learning rate (0.002) was reduced by a factor of 10 when the validation loss did not improve for 10 consecutive epochs.

The output of ScoreCAM was binarized with the threshold of 0.05 and used as pseudo-labels for the U-Net model, which was subsequently trained to segment the artifacts using the same datasets’ splits and training procedure. All the experiments were conducted using a Tesla V100-PCIE-32GB Graphics Processing Unit.

#### 2.3.2. Comparison with other methods

We compared the segmentation results obtained from the ScoreCAM-U-Net to a number of alternative solutions. As ScoreCAM-U-Net is a combination of ScoreCAM and U-Net models, we first compared our performance to each of these models separately. We expected a strongly supervised U-Net model trained on pixel-level annotations to show better performance than its weakly supervised counterparts. We also compared the proposed approach to the current state-of-the-art algorithms used to detect anomalies using image labels in domains other than microscopy: Patch Support Vector Data Description (PatchSVDD)^26,27^, Patch Distribution Modeling (PaDiM)^27^, and an autoencoder-based method (AE)^24^. All model architectures, training parameters and training processes are adopted here as defined in the original papers^24,26,27^.

PaDiM and PatchSVDD are both embedding similarity-based methods that use convolutional neural network-based approaches (encoders) that learn robust and short representations (embeddings) from patches of clean images. During the inference, the encoders are used to extract embeddings from test image patches and compare them to the embeddings extracted from the clean images based on a similarity metric. The main difference between these two methods is in the similarity metric employed to compare the embeddings as well as in the way the embeddings are constructed. PaDiM applies the Mahalanobis distance metric^39^ and constructs an embedding by combining the features of multiple encoder layers, whereas PatchSVDD uses the Euclidean distance metric and constructs the representation from the features of a single encoder layer. Based on the embedding comparisons, each test image patch is assigned a similarity score in which a low similarity score indicates the presence of artifacts. The final segmentation of each test image is constructed after the similarity scores of these patches are distributed to their pixels and the corresponding patch segmentations are merged together.

The AE method also utilizes a convolutional neuronal network based approach (an encoder-decoder network architecture) that first learns representations of the clean input images (using the encoder) and then to reconstruct the original clean input images from the learned representations (via the decoder). During inference, the trained model is expected to fail to reconstruct the artifactual areas of the test images as the network has only acquired rich representations of clean images. Therefore, artifacts manifest themselves in areas with a high pixel-wise difference between the input image and its reconstructed counterpart.

We measure the ability of the models to correctly identify the presence of an artifact in the image using the F1 score which is the harmonic mean of precision and recall. We also assess the segmentation performance via calculating pixel-wise precision, recall, F1, and the intersection over union (Box 1).

##### Box 1. Performance measures

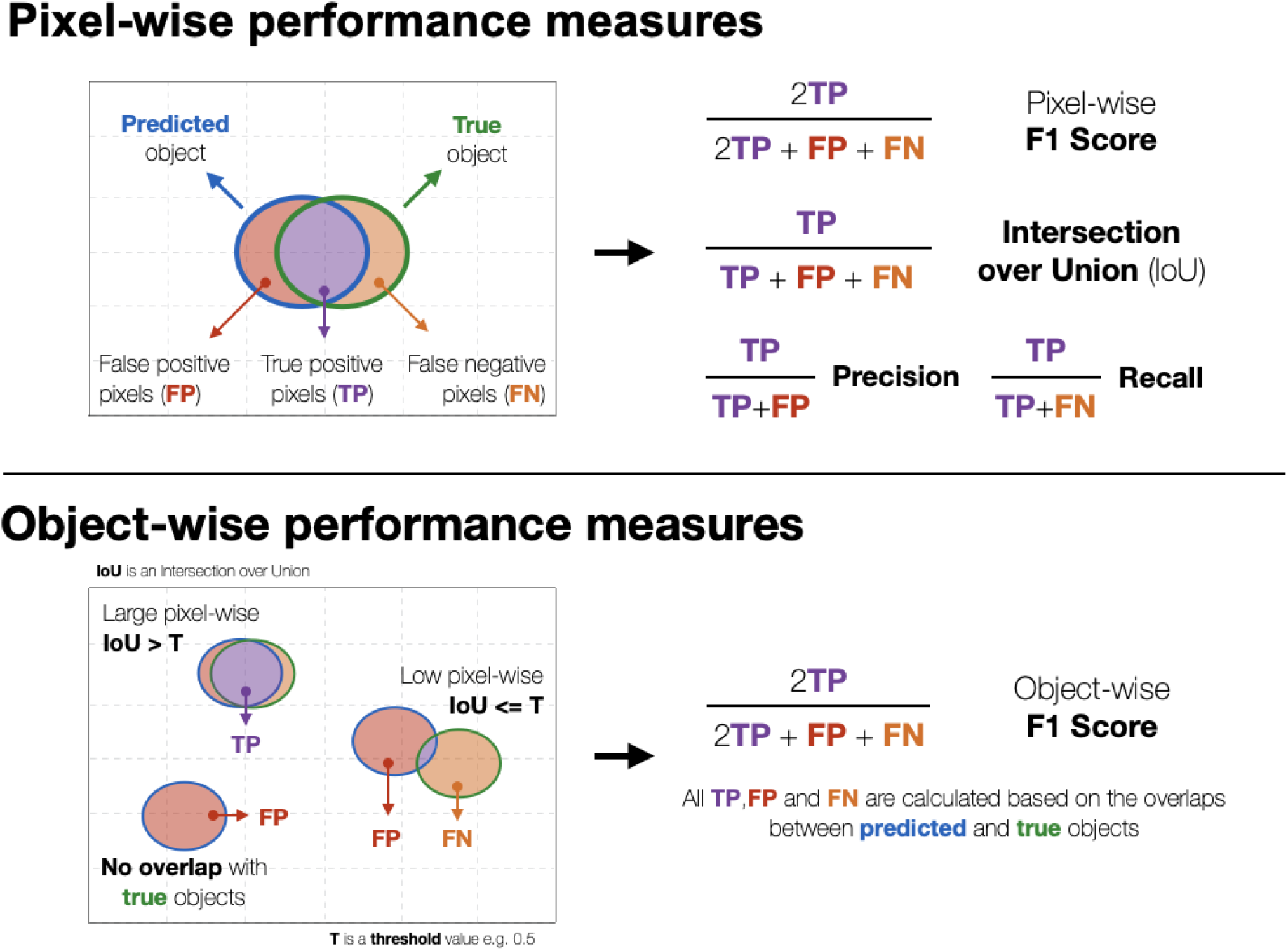

#### 2.3.3. Post-processing

We first binarized the probability maps produced by the models at cutoffs of 0.75 for AE, 0.3 for PaDIM, 0.0005 for PatchSVDD, 0.001 for ScoreCAM, 0.001 for ScoreCAM-U-Net and 0.45 for U-Net. These cutoffs were selected to maximize pixel-wise IoU (Box 1) performance on validation data. We then filtered out objects smaller than 1000, 500, and 500 pixels in the seven cell lines, the ArtSeg-CHO-M4R, and the LNCaP datasets respectively using *remove_small_objects* function from the skimage package^40^. The sizes of the filtered-out objects were selected to maximize the pixel-wise IoU of the majority number of models, and different sizes do not drastically change the performance of the models (Supplementary Tables 2, 3, and 4). We recommend using expert knowledge to select the size of objects to filter out.

#### 2.3.4. Measuring impact of artifacts and artifact removal on the downstream analyses

To evaluate the utility of removing artifacts in microscopy experiments, we focused on two common types of downstream analyses: nuclei segmentation and effective concentration estimation from concentration-response assays. The former is a standard step in the majority of cell microscopy workflows while the latter is an example of a commonly used pipeline where cell segmentation is used for image intensity quantification which is followed up by regression analysis.

##### Nuclei segmentation

In order to assess how nuclei segmentation accuracy inside the artifactual regions compares to artifact-free areas, we evaluated the performance of nuclei segmentation in the seven cell lines dataset inside and outside the artifactual areas. To detect and segment the nuclei from the brightfield images we used an existing PPU-Net^3^ model. The training, ground truth preparation, and post-processing steps for this model are described in the original publication^3^. We calculated segmentation pixel-wise F1 and object-wise F1 scores (Box 1), following previously described approaches^3,41^, and morphological properties (size and solidity) of the resulting nuclei.

##### Ligand affinity estimation

In downstream analysis of pharmacological experiments, the cell bodies are segmented from brightfield images using a U-Net-based deep learning model^28^, and the cell fluorescence intensities are quantified from a parallel fluorescence channel based on the segmentation. The fluorescence intensities of cells depend on the strength of interaction (affinity) between the protein and the interaction partner (ligand) as well as the ligand concentration. The strength of protein-ligand interaction is determined using regression analysis of competitive ligand concentrations and the well average fluorescence intensity information from up to 64 individual images.

We studied the impact of artifacts and artifact removal on the determination of receptor-ligand interaction affinity. For that, in each of the ten individual concentration-response experiments, the cells were detected from brightfield images using a previously developed U-Net based segmentation model with an F1-score of 0.89^28^. The artifactual areas determined manually or with ScoreCAM-U-Net were removed from the analysis. For experimental control, the analysis was also carried out without any artifact removal. The average intensity of the detected cell pixels as well as the average intensity of the background were determined from the aligned red fluorescence protein filter (excitation: 531(40) nm, emission: 593(40) nm) fluorescence images made in parallel with the brightfield images. The values were averaged for all images from the same well. For each well, to find the specific fluorescence intensity of bound fluorescence ligand the difference between cellular and background fluorescence intensities was calculated. LogIC50 values corresponding to half maximal displacement of the fluorescence ligand were obtained via nonlinear regression analysis. For that, the fluorescence intensity dependence on the competitive ligand concentration was fitted with the Hill equation using GraphPad Prism 5.0 and “log(inhibitor) vs. response” nonlinear regression model which is equivalent to the logistic regression.

Concentration-response experiments serve as a good example for image analysis pipelines that rely on image intensity calculation and regression in the downstream analysis. For quantifying the quality of the full pipeline, we chose the absolute difference between the LogIC_50_ values calculated from manual artifact removal and the alternative option. The difference of LogIC_50_ values describes how accurate pharmacological parameters can be obtained with and without anomaly removal. We also used the R^2^ value of the Hill equation fit as a metric, which reflects the overall agreement between the experiment and the model. Finally, we chose the Pearson’s correlation coefficient *r* between predicted fluorescence intensity values using manual artifact removal and the alternative method, which allows isolating the effect of artifacts on the signal directly without the influence of other sources of uncertainty.

## 3. Results

To develop and test a weakly supervised method for artifact segmentation: we confirmed that artifacts exist and are prevalent in brightfield microscopy images; annotated artifacts in three datasets; tested models for finding them automatically; and evaluated the impact of removal on downstream analysis results.

### 3.1. Artifacts in brightfield images are prevalent and diverse

The artifacts in the seven cell lines dataset range from very big (e.g. a clump of detached cells covering 49% of the image pixels) to tiny ones only a few pixels in size. The average annotated artifact size in this dataset is 4,417 pixels, which is larger than a typical nucleus in this dataset, and 16% of images had at least 10% of their area covered by artifacts. The artifacts in the seven cell lines dataset were heterogeneous in their size and morphological properties (Figure 2, Supplementary Figure 2).

**Figure 2:**
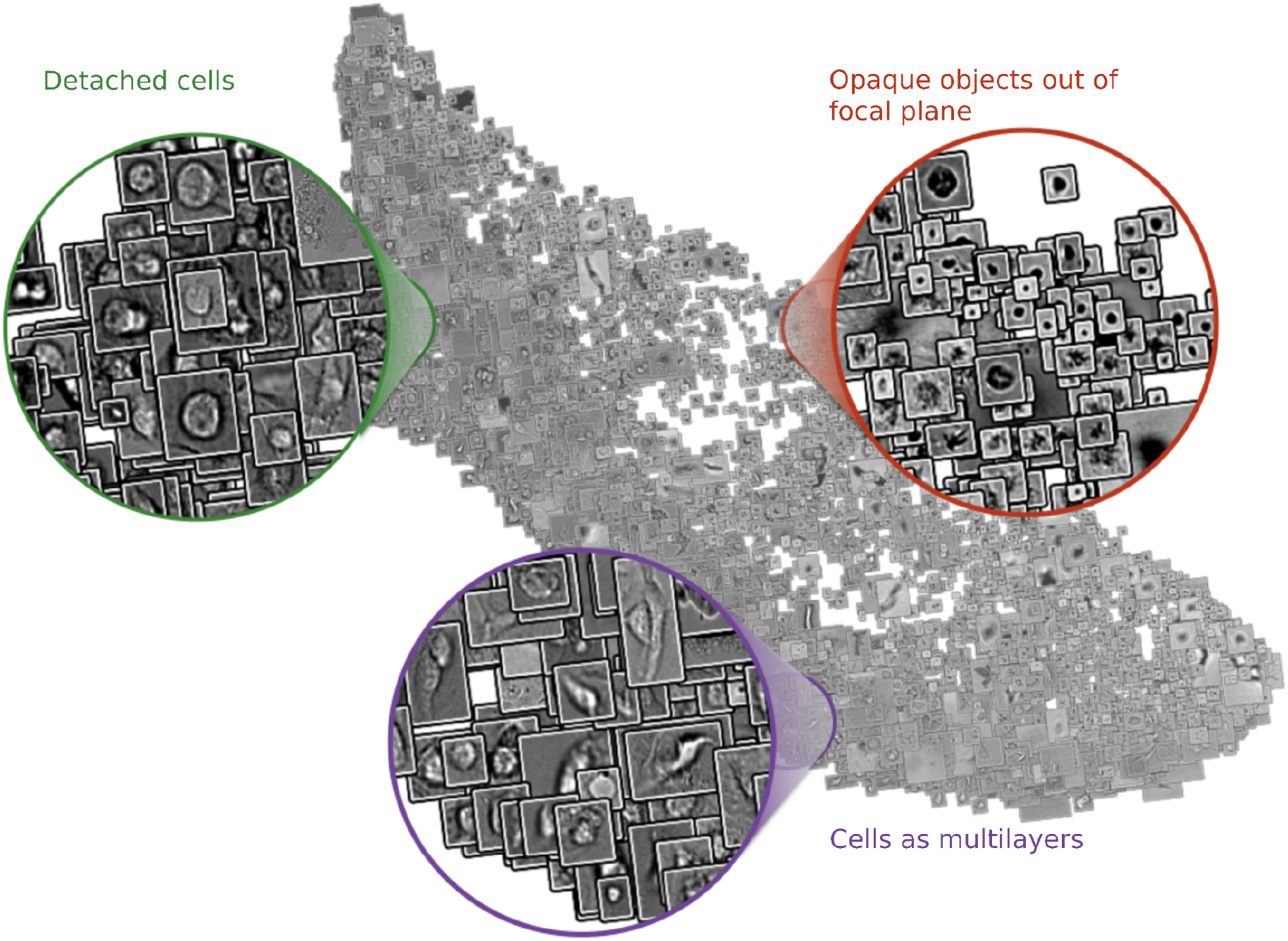
Artifacts are heterogeneous, and range in shapes and sizes. A UMAP projection of all artifacts from the seven cell lines dataset. The inputs to the UMAP are the pixels of each patch that contains an artifact and the outputs are the first two features in the UMAP embeddings of each patch. We then used these two features respectively as ‘x’ and ‘y’ values to plot the corresponding input patch in 2D space.

In the LNCaP dataset, we annotated 60 objects that affected 6.5% of the images. The sizes of artifacts range from big (e.g. a hair covering 10% of the image pixels) to small, which covers only 0.07 % of the pixels, with the average artifact being 75,933 pixels (Supplementary Figure 2).

In the ArtSeg-CHO-M4R dataset, almost all images had artifacts, with a total of 13,713 artifact objects in 1,171 affected images. Again, the largest object covered a large part of the image (e.g. 63% as a clump of detached cells), while the smallest one was a few pixels in size (Supplementary Figure 2). An average artifact in this dataset had an area of 3,450 pixels, or 0.31% of image size.

### 3.2. Artifacts can be accurately detected with weak supervision

Next, we compared different approaches for artifact detection and segmentation qualitatively and quantitatively (Figure 3 A,B; Supplementary Figure 3). We first evaluated the ability of the models to detect artifacts in the images. As ScoreCAM-U-Net and ScoreCAM both use the same ResNet classification backend, their detection performance is the same, with both models achieving image classification F1 scores of 93.2%, 93.7% and 90% in seven cell lines, LNCaP and ArtSeg-CHO-M4R datasets respectively (Figure 3B, Supplementary Table 1). Other methods were less accurate, with the only exception of U-Net outperforming ScoreCAM-based models in the LNCaP dataset (99.4% F1 score for U-Net over 93.7% for ScoreCAM-U-Net; Figure 3B).

**Figure 3:**
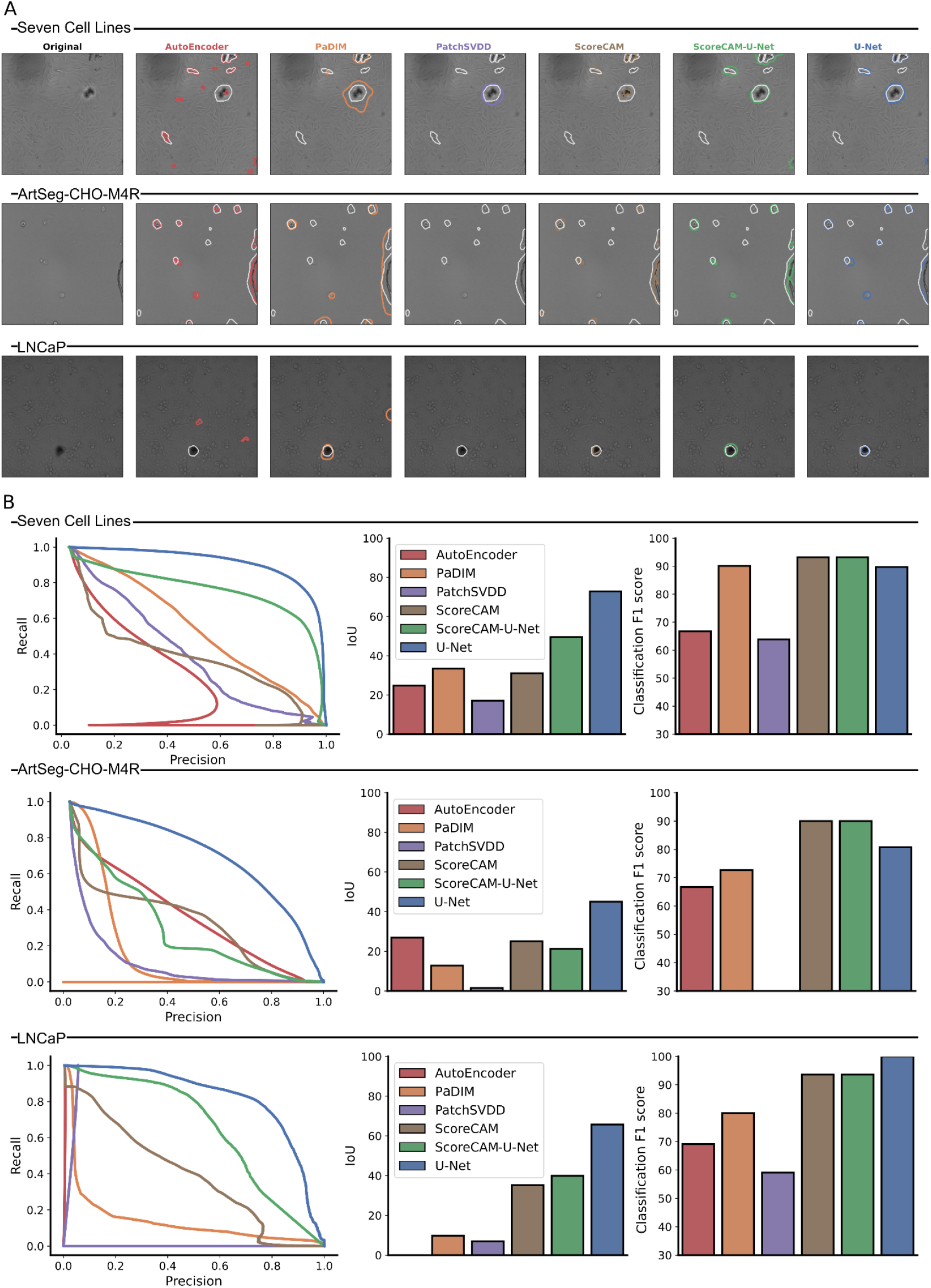
Artifact segmentation and image-level classification results for all models (colors) in seven cell lines, LNCaP, and ArtSeg-CHO-M4R datasets. **A)** Examples of brightfield images and the corresponding artifact segmentation of all models (columns, colors) and datasets (rows; separated by lines and dataset names). White contour: expert annotated artifact boundaries; colored contours: artifact segmentation boundaries of the corresponding model. **B)** Different performance metrics for all models (colors) and datasets (rows). Left column: artifact segmentation precision (x-axis) and recall (y-axis) of artifact detection at different thresholds (points along the curve) for all models and datasets. Middle column: artifact segmentation pixel-wise IoU (y-axis) for all models and all datasets. Right column: image-level classification F1 score (y-axis) for all models (x-axis) and datasets (rows).

We then assessed the models’ performance in segmenting artifacts. ScoreCAM-U-Net outperforms the other non-strongly supervised models by achieving the highest area under the precision-recall curve, as well as the largest average object intersection over union on seven cell lines and LNCAP datasets (Figure 3B). There was no dominant weakly supervised model in the ArtSeg-CHO-M4R dataset. Compared to the strongly supervised U-Net model, ScoreCAM-U-Net got the second-highest IoU performance in the seven cell lines (49.5 ScoreCAM-U-Net vs 72.9 U-Net) and the LNCaP (39.9 ScoreCAM-U-Net vs 65.74 U-Net) datasets (Supplementary Table 1).

Although the strongly supervised approach outperformed weakly supervised methods, it took substantial time to prepare the pixel-level annotations required for the U-Net model compared to weak labeling. On average, an expert spent 279 seconds to produce pixel-level annotation for a single microscopy image, while it took them only 2 seconds to point out if a given image contained an artifact. Hence, weakly supervised methods consume about two orders of magnitude less of expert time for the given case. Therefore, when making a choice of method for dealing with artifacts, it is reasonable to take into account the dataset size and the amount of time needed to produce relevant annotations. For complex datasets which require large training datasets for model development, generating precise pixel-level labels would be very time consuming, and hence it is practical to prefer weakly supervised approaches like ScoreCAM-U-Net.

### 3.3. Weakly supervised artifact removal improves downstream analysis

After establishing the quality of the proposed ScoreCAM-U-Net method for artifact detection and segmentation, we evaluated the impact of using it for cleaning images on two downstream applications.

#### 3.3.1. Removing artifacts improves quality of nuclei segmentation

As artifacts distort pixels that otherwise represent nuclei (Figure 4), we observed substantial degradation in nuclei segmentation performance due to artifacts. The pixel-wise F1 score decreased from 0.89 in artifact-free to 0.60 in artifactual regions; and the object-wise F1 score decreased from 0.65 in artifact-free to 0.28 in artifactual regions (Figure 5). This had a direct impact on naive analyses that do not differentiate between artifactual and clean regions, reducing segmentation accuracy (0.87 pixel-wise F1, 0.61 object-wise F1; Figure 5). Importantly, automatically removing artifacts using ScoreCAM-U-Net has the same impact as manual removal, improving the segmentation performance to near-optimal 0.89 and 0.64 pixel-wise and object-wise F1 scores (Figure 5).

**Figure 4:**
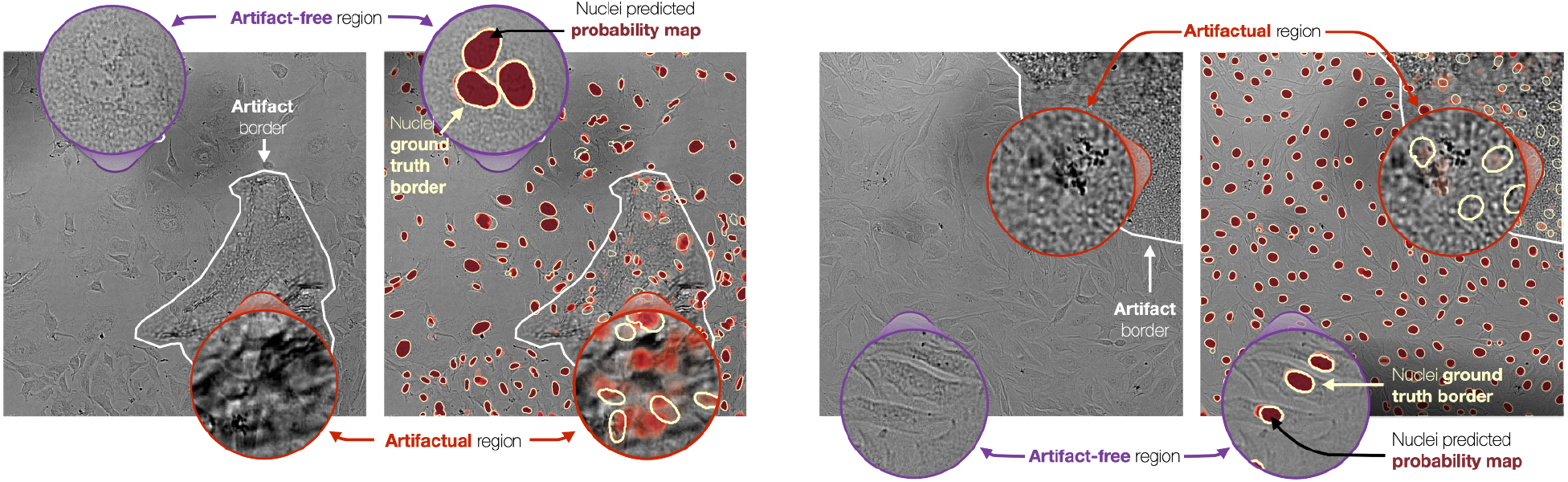
Visual impact of artifacts on nuclei segmentation. Two pairs of brightfield images with corresponding nuclei segmentation (in dark red) overlaid. Zoomed-in purple circles represent examples of artifact-free areas and artifactual areas (light red). White contours: artifact borders; yellow-ish white contours: nucleus ground truth borders; arrows and text: guides to corresponding regions and elements.

**Figure 5:**
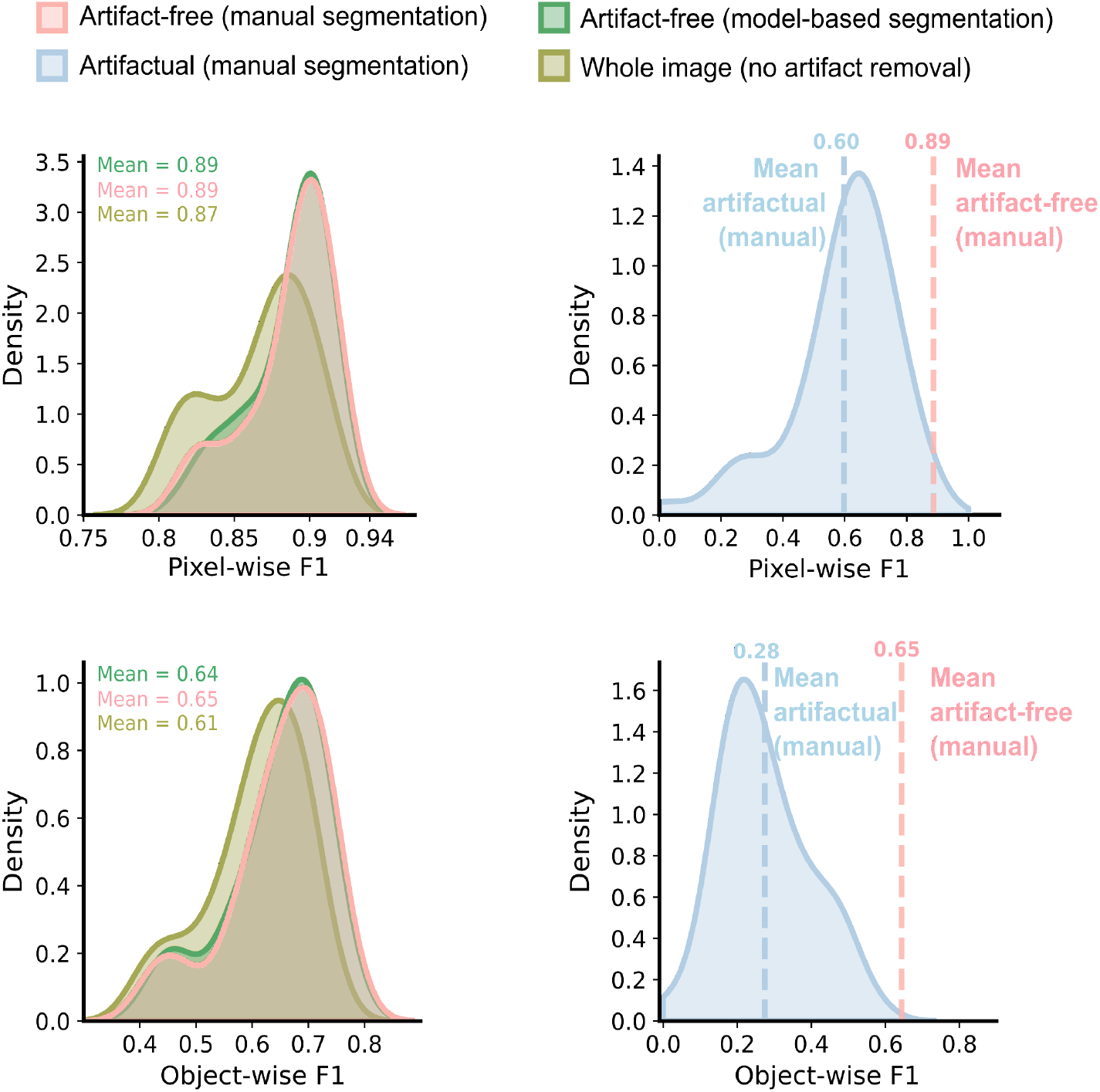
Impact of artifacts and artifact removal on downstream analyses. Density (y-axis) of image-average nucleus segmentation pixel-wise F1 (top, x-axis) and object-wise F1 (bottom, x-axis) in the seven cell lines dataset for different areas of the image (colors). Pink: area in the images manually annotated as not artifacts; blue: area in the images manually annotated as artifacts; green: area in the images automatically annotated as not artifacts by ScoreCAM-U-Net; yellow: all image area. Dashed lines: mean pixel-wise F1 and object-wise F1 of segmented nuclei in the artifactual and artifact-free regions (different colors).

We next considered nuclear size and morphology metrics with and without artifact correction. Nuclei in areas containing artifacts show different morphological properties with nuclei solidity of 0.92 and size of 213 pixels while the same properties are 0.95 and 400 pixels respectively in the artifact-free regions (Supplementary Figure 4). In concordance with the segmentation results, automatically removing artifacts using ScoreCAM-U-Net recovers the expected nuclei size and solidity of 397 pixels and 0.95 respectively for artifact-free areas, again performing close to the gold standard of manual removal (Supplementary Figure 4). These results demonstrate that automatic removal can overcome the detrimental effect of artifacts with quality close to manual filtering.

#### 3.3.2. Removing artifacts improves pharmacological parameter estimates

Cell segmentation in images is a commonly used process to determine biochemical or pharmacological parameters from microscopy experiments. Common examples of such experiments include quantifying image intensity of the segmented areas. This can be followed up by a test of significance or regression analysis to determine biochemical parameters like half-life of a reaction or the half maximal effective concentration of a substance. We analyzed how presence of artifacts affects the quality of a microscopy image-based analysis used to determine ligand affinity to M_4_ muscarinic receptors using the ArtSeg-CHO-M4R dataset. Manual anomaly removal has a clear effect on both the plateau locations and the estimated Log(IC_50_) values (Figure 6, Supplementary Figure 5). The mean absolute difference between Log(IC_50_) calculated with manual artifact removal and no artifact removal is 0.29 units, equivalent to a two-fold error in dose. In contrast, after automatic artifact removal with ScoreCAM-U-Net, the Log(IC_50_) difference from manual anomaly removal was reduced to just 0.16 units, which is similar to the standard deviation of 0.11 observed between biological replicate experiments. The model fit explained 0.89, 0.86, and 0.74 of the data variation for manual anomaly removal, ScoreCAM-U-Net anomaly removal, and no anomaly removal respectively. Finally, the Pearson’s correlation coefficient of well-average fluorescence intensities between manual anomaly removal and ScoreCAM-U-Net anomaly removal is 0.98 while the correlation between manual anomaly removal and no anomaly removal is only 0.93. Overall, removing artifacts leads to an increase in replicate correlation, which itself results in reduction in estimate uncertainty. The estimated ligand affinity better reflects the values established from manually cleaned images. This confirms that artifact removal leads to considerable improvement of downstream regression or statistical analysis which relies on image intensity quantification.

**Figure 6:**
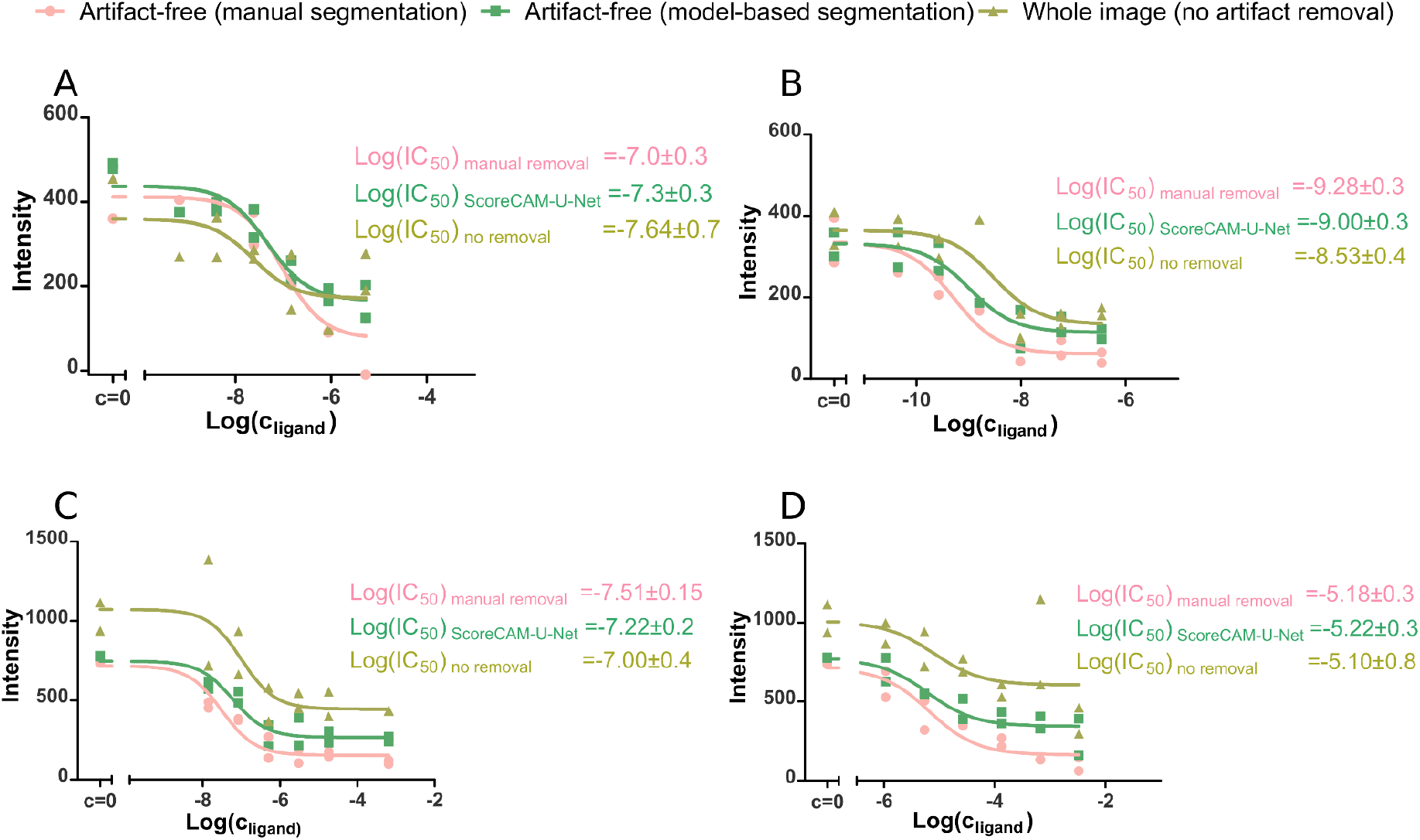
Cell fluorescence intensity dependence on M4 receptor ligand concentration determined with live-cell fluorescence microscopy at the presence of 2 nM UR-CG072. Displacement curves of three different ligands are shown: pirenzepine (**A** and **C**), atropine (**B**) and arecholine (**D**). Three different artifact removal methods at the image analysis stage are compared (colors): manual artifact segmentation, ScoreCAM-U-Net segmentation and no artifact removal. For each combination of ligand and artifact removal method a regression analysis is performed with Hill equation (Hill coefficient fixed at −1) with the best fits shown as continuous lines. For each displacement curve, the Log(IC_50_)±SD is presented, where SD represents the standard deviation estimation of Log(IC_50_). Each displacement curve was measured in duplicates with each data point representing the average fluorescence intensity of cells in each well.

## 4. Discussion

We proposed ScoreCAM-U-Net, a deep learning model for identifying artifacts in brightfield microscopy images that combines the benefits of weakly supervised learning which does not require delineating objects, and strongly supervised learning that provides pixel-level resolution. As training is performed using only image-level labels, generating training data is orders of magnitude faster, but without substantially sacrificing performance compared to pixel-level annotation. To our knowledge, this is the first attempt to automatically detect artifacts in large sets of brightfield microscopy images.

Our results demonstrate that artifacts have an adverse impact on nuclei segmentation and that detection and measurement of nuclei are improved when removing such artifacts. We showed that this impact manifests in both quantitative segmentation metrics such as pixel-wise and object-wise F1 score, as well as morphological properties of the nuclei like solidity and size, which are central for cytometry applications. Almost all study designs that use large-scale cell microscopy and image quantification-based readout would benefit from our model.

One important application of cell microscopy is intensity quantification for studying the localization and co-localization of fluorescently labeled molecules. To exemplify this type of analysis, we studied how artifact removal affects the calculation of drug-receptor binding affinities based on live-cell fluorescence and brightfield microscopy. After artifact removal with ScoreCAM-U-Net, the estimated ligand affinities are in better agreement with the values established from manually cleaned images. The model-based estimates also reduce linear regression uncertainty and result variability of independent experiments, indicating a combination of better fit of the theoretical model and improved reproducibility of the measurements. Thus, artifact removal improves image intensity quantification independent of the nature of statistical analysis applied downstream.

Our ScoreCAM-U-Net method establishes the utility of automatically segmenting artifacts from brightfield microscopy images. The key benefit of our approach is its scalability, such that clean images can be obtained for screening campaigns that would be unreasonable to process manually, while its downside is an inability to differentiate different types of artifacts. For example, the current model would not tell if an image contains an artifact of cell debris and the other contains bacterial contamination. A natural extension can build on our approach to train a model that can differentiate between different types of artifacts. Other extensions can use the power of deep learning for other imaging modalities, such as histopathology, as well as to further reduce annotation time. We envision that ultimately, all common artifacts will be automatically segmented and optionally removed at the time of acquisition with no input needed by the operator. Moreover, we believe that the encouraging results presented in this work will motivate use of weakly supervised segmentation methods such ScoreCAM-U-Net in other areas where pixel-level annotations are prohibitively expensive or time-consuming to acquire, i.e. medicine.

## Acknowledgments

MA and KH were supported by the Estonian Research Council (PRG1095, PSG59) and the Estonian Centre of Excellence in IT (EXCITE) (TK148). TL, JT, MT, and AR were supported by the University of Tartu ASTRA Project PER ASPERA, financed by the European Regional Development Fund, COST action CA 18133 ERNEST, and the Research Training Group GRK1910 of the Deutsche Forschungsgemeinschaft (DFG). LP was supported by Wellcome (206194), the Estonian Research Council (IUT34-4), and the Estonian Centre of Excellence in IT (EXCITE) (TK148). KP was supported by PerkinElmer Cellular Technologies. DF was supported by Estonian Research Council grants (PRG1095, PSG59 and ERA-NET TRANSCAN-2 (BioEndoCar)); Project No 2014-2020.4.01.16-0271, ELIXIR and the European Regional Development Fund through EXCITE Center of Excellence.

We thank the High-Performance Computing center at the University of Tartu, Institute of Computer Science for providing the computational resources. We thank Dr. Max Keller from the University of Regensburg for providing ligands UNSW-MK259, UR-SK75 and UR-CG072. We thank Erika Charola for proofreading and language editing.

## Author contributions

Designed project: DF, LP, MA, KH, TL. Performed experiments: KH, MA, TL. Performed analysis: MA, KH, TL, DF. Supervised study: DF, LP, KP, AR. Prepared data: MT, JT, TL, KH. Wrote paper: MA, TL, LP, DF with input from all authors.

## Data availability

The ArtSeg-CHO-M4R dataset is publicly available at https://datadoi.ee/handle/33/433; with doi: http://dx.doi.org/10.23673/re-307.

## Supplementary Material

**Supplementary Table 1:** Segmentation and image-level detection/classification results in seven cell lines, ArtSeg-CHO-M4R, and LNCaP datasets. PW: pixel-wise; IoU: intersection over union.

| Dataset | Metric | U-Net | ScoreCAM-U-Net | ScoreCAM | AE | PADIM | Patch_SVDD |
| --- | --- | --- | --- | --- | --- | --- | --- |
| Seven Cell Lines | Classification F1-Score | 89.7 | 93.2 | 93.2 | 66.7 | 90.1 | 63.9 |
|  | Segmentation PW-F1 | 84.4 | 66.2 | 47.2 | 39.7 | 50 | 28.3 |
|  | Segmentation PW-precision | 86 | 74.4 | 42 | 32.4 | 59.7 | 39.6 |
|  | Segmentation PW-recall | 82.8 | 61 | 54.6 | 51.4 | 43.1 | 23.9 |
|  | Segmentation IoU | 72.9 | 49.5 | 31.1 | 24.8 | 33.4 | 17.1 |
| ArtSeg-CH O-M4R | Classification F1-Score | 80.8 | 90 | 90 | 66.7 | 72.7 | 23.6 |
|  | Segmentation PW-F1 | 62 | 34.9 | 40.1 | 42.4 | 22.6 | 3.1 |
|  | Segmentation PW-precision | 79 | 30.3 | 58.1 | 47.3 | 36.1 | 40.4 |
|  | Segmentation PW-recall | 51.4 | 41.5 | 30.9 | 38.5 | 19.4 | 26.6 |
|  | Segmentation IoU | 45 | 21.3 | 25.1 | 26.9 | 12.8 | 1.6 |
| LNCaP | Classification F1-Score | 99.4 | 93.7 | 93.7 | 69.1 | 80 | 59.1 |
|  | Segmentation PW-F1 | 79.3 | 55.7 | 52 | 0 | 17.7 | 11.6 |
|  | Segmentation PW-precision | 75 | 50.5 | 75.2 | 0 | 17.8 | 9.9 |
|  | Segmentation PW-recall | 84.4 | 79.5 | 40.9 | 0 | 17.6 | 41.6 |
|  | Segmentation IoU | 65.7 | 39.9 | 35.2 | 0 | 9.7 | 6.9 |

**Supplementary Figure 1:**
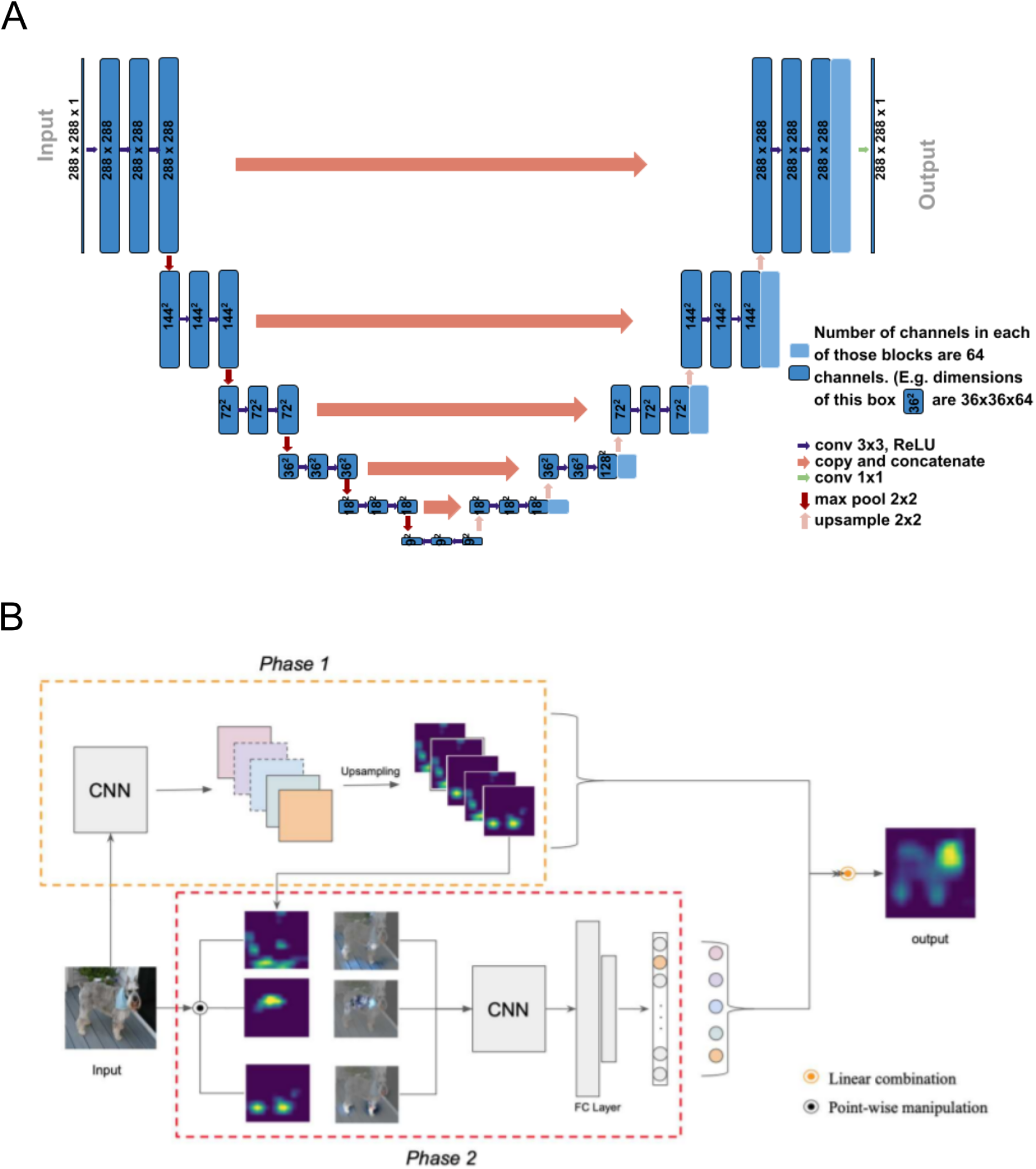
Zoomed components of the pipeline. **A**) U-Net model used for object segmentation. **B**) ScoreCAM^18^ algorithm which is used to generate the ground truth to train U-Net during training, and used to determine whether there are any artifact objects in the image during inference.

**Supplementary Table 2:** Intersection over union of all models in the seven cell lines dataset after removing predicted objects smaller than different cutoff sizes

| **Object size cutoff(pixels)** | **AE** | **PaDIM** | **Patch_SVDD** | **ScoreCAM** | **ScoreCAM-U-Net** | **U-Net** |
| --- | --- | --- | --- | --- | --- | --- |
| 20 | 21.1 | 33.4 | 27.9 | 23.4 | 52 | 72.9 |
| 50 | 21.2 | 33.4 | 27.9 | 23.5 | 52.1 | 72.9 |
| 100 | 21.5 | 33.4 | 27.9 | 23.5 | 52.1 | 72.9 |
| 500 | 24.8 | 33.4 | 27.9 | 23.9 | 53.1 | 72.9 |
| 1000 | 28.6 | 33.3 | 27.9 | 24.2 | 53.6 | 73 |
| 2000 | 33.3 | 33.2 | 27.9 | 24.2 | 54.1 | 73.2 |

**Supplementary Table 3:** Intersection over union of all models in the ArtSeg-CHO-M4R dataset after removing predicted objects smaller than different cutoff sizes

| **Object size cutoff (pixels)** | **AE** | **PaDIM** | **Patch_SVDD** | **ScoreCAM** | **ScoreCAM-U-Net** | **U-Net** |
| --- | --- | --- | --- | --- | --- | --- |
| 20 | 26.8 | 13.4 | 3.4 | 27.3 | 23.1 | 44.4 |
| 50 | 26.8 | 13.4 | 3.4 | 27.3 | 23.1 | 44.4 |
| 100 | 26.9 | 13.4 | 3.4 | 27.3 | 23.1 | 44.4 |
| 500 | 26.9 | 13.3 | 3.4 | 26.5 | 22.9 | 43.7 |
| 1000 | 24.2 | 13.1 | 3.4 | 24.8 | 22.3 | 42 |
| 2000 | 18 | 12.3 | 3.4 | 20.3 | 20.2 | 35 |

**Supplementary Table 4:** Intersection over union of all models in the LNCaP dataset after removing predicted objects smaller than different cutoff sizes

| **Object size cutoff (pixels)** | **AE** | **PaDIM** | **Patch_SVDD** | **ScoreCAM** | **ScoreCAM-U-Net** | **U-Net** |
| --- | --- | --- | --- | --- | --- | --- |
| 20 | 0 | 9.6 | 2.8 | 28.2 | 44.8 | 62.9 |
| 50 | 0 | 9.6 | 2.8 | 28.3 | 44.8 | 62.9 |
| 100 | 0 | 9.6 | 2.8 | 28.3 | 44.8 | 62.9 |
| 500 | 0 | 9.7 | 2.8 | 29.1 | 44.7 | 62.9 |
| 1000 | 0 | 9.3 | 2.8 | 28.5 | 44.7 | 62.9 |
| 2000 | 0 | 9.6 | 2.8 | 29.4 | 44.7 | 63.8 |

**Supplementary Figure 2:**
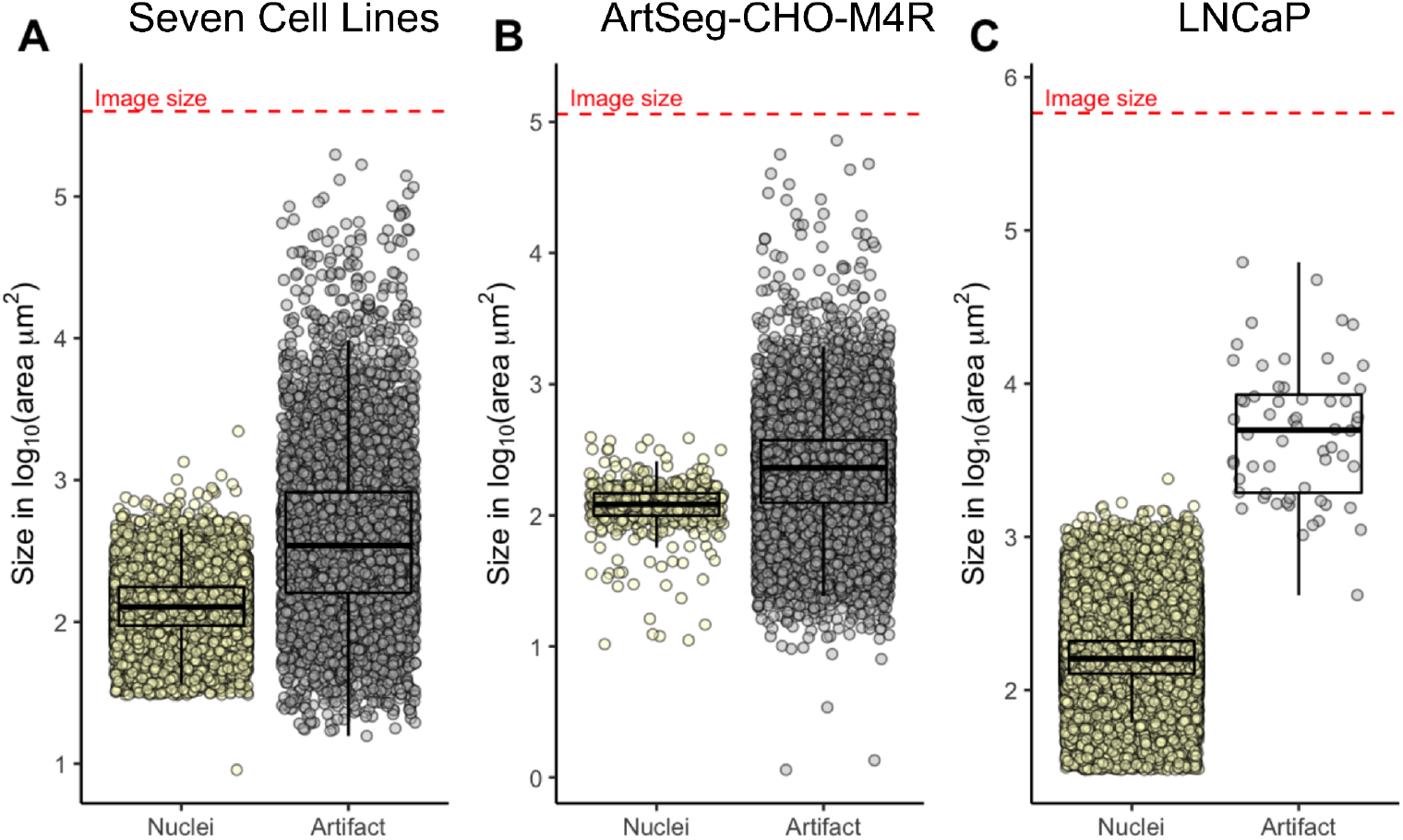
Artifacts are in different sizes. Size (mm^2^ in log_10_ scale, y-axis) of nuclei and artifact objects (colors, x-axis) in seven cell lines **(A)**, ArtSeg-CHO-M4R **(B)**, and LNCaP datasets **(C)**. Boxes: 25th, 50th and 75th percentile; whiskers: 1.5x from the interquartile range; circles: individual artifact or nucleus. Dashed red line: size of the whole image.

**Supplementary Figure 3:**
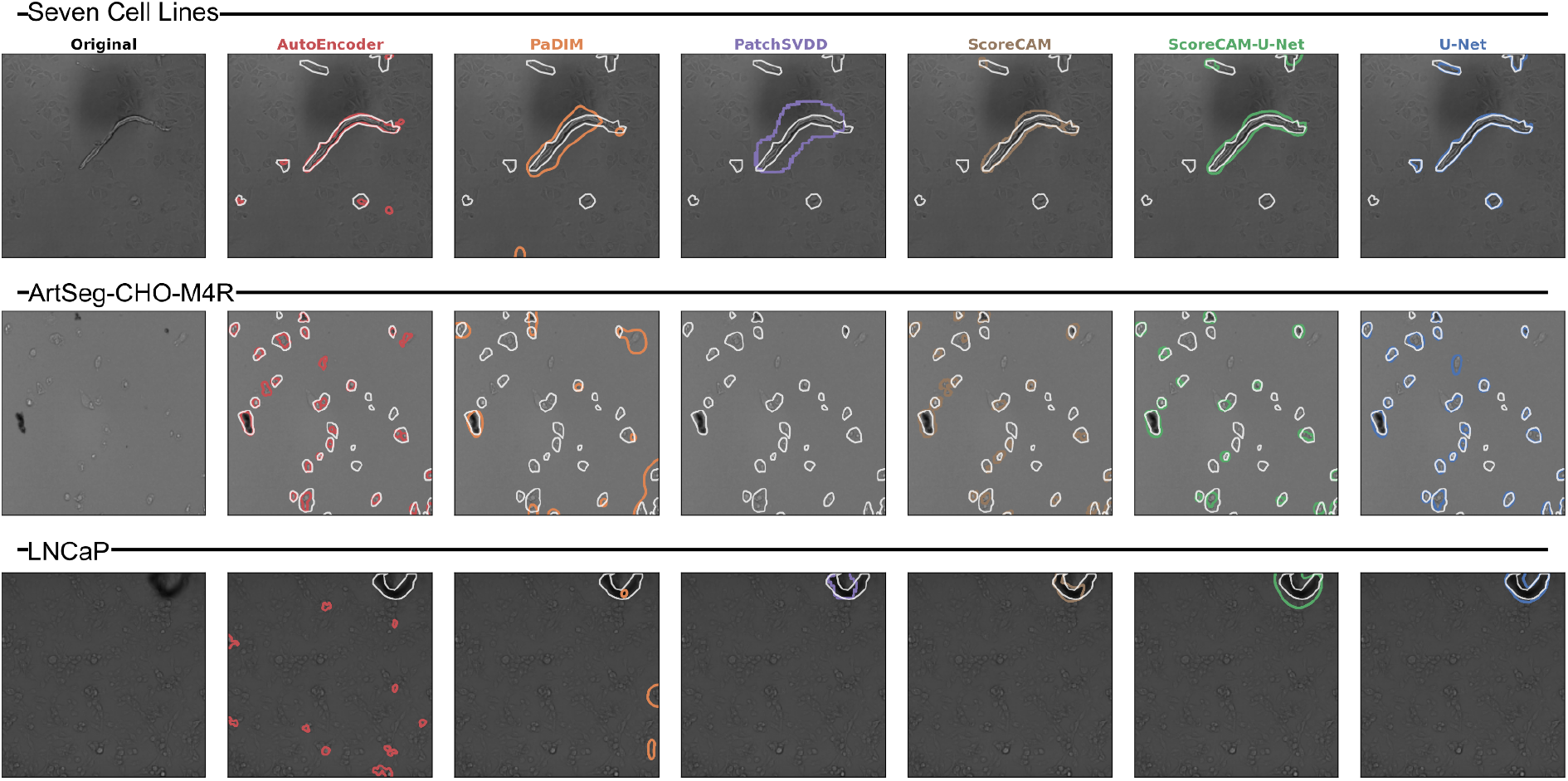
Artifact segmentation examples for all models (colors) in seven cell lines, LNCaP, and ArtSeg-CHO-M4R datasets (rows). Examples of brightfield images and the corresponding artifact segmentation of all models (columns, colors) and datasets (rows; separated by lines and dataset names). White contour: true artifact boundaries; colored contours: artifact segmentation boundaries of the corresponding model.

**Supplementary Figure 4:**
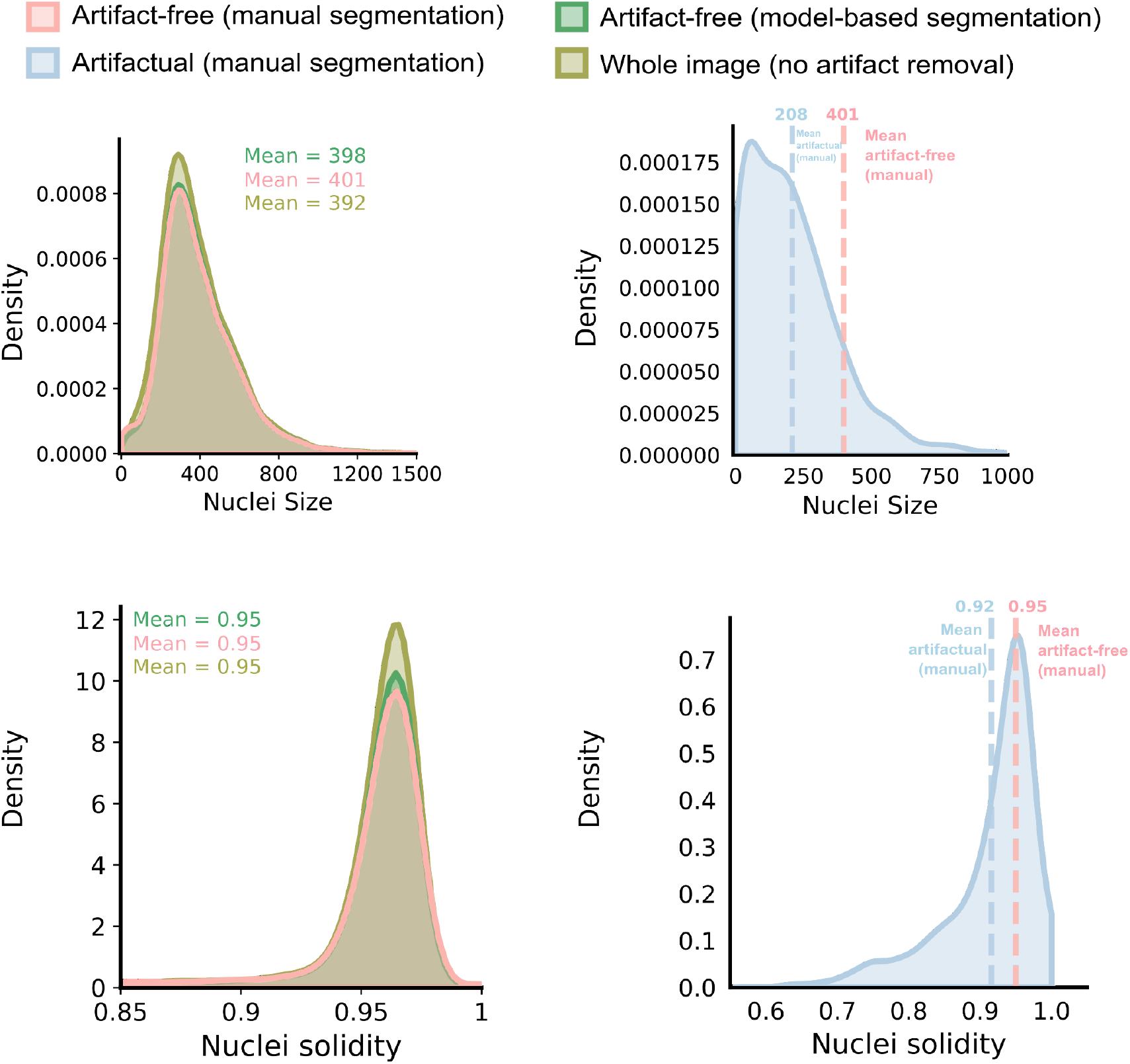
Impact of artifacts and artifact removal on downstream analyses. Density (y-axis) plots of segmented nucleus size in pixels (top, x-axis) and solidity (bottom, x-axis) in the seven cell lines dataset for different areas of the image (colors). Metrics are calculated for different areas of the images (colors); artifactual: artifactual areas in the images; artifact-free: area in the images other than the artifactual areas. Artifacts are detected in two ways; manual: the detection of artifacts is performed manually; model-based: the ScoreCAM-U-Net model is used to detect the artifacts. No artifact removal: the metrics are calculated without removing artifacts. Dashed lines: mean size in pixels and solidity of segmented nucleus in the artifactual and artifact-free regions(different colors).

**Supplementary Figure 5:**
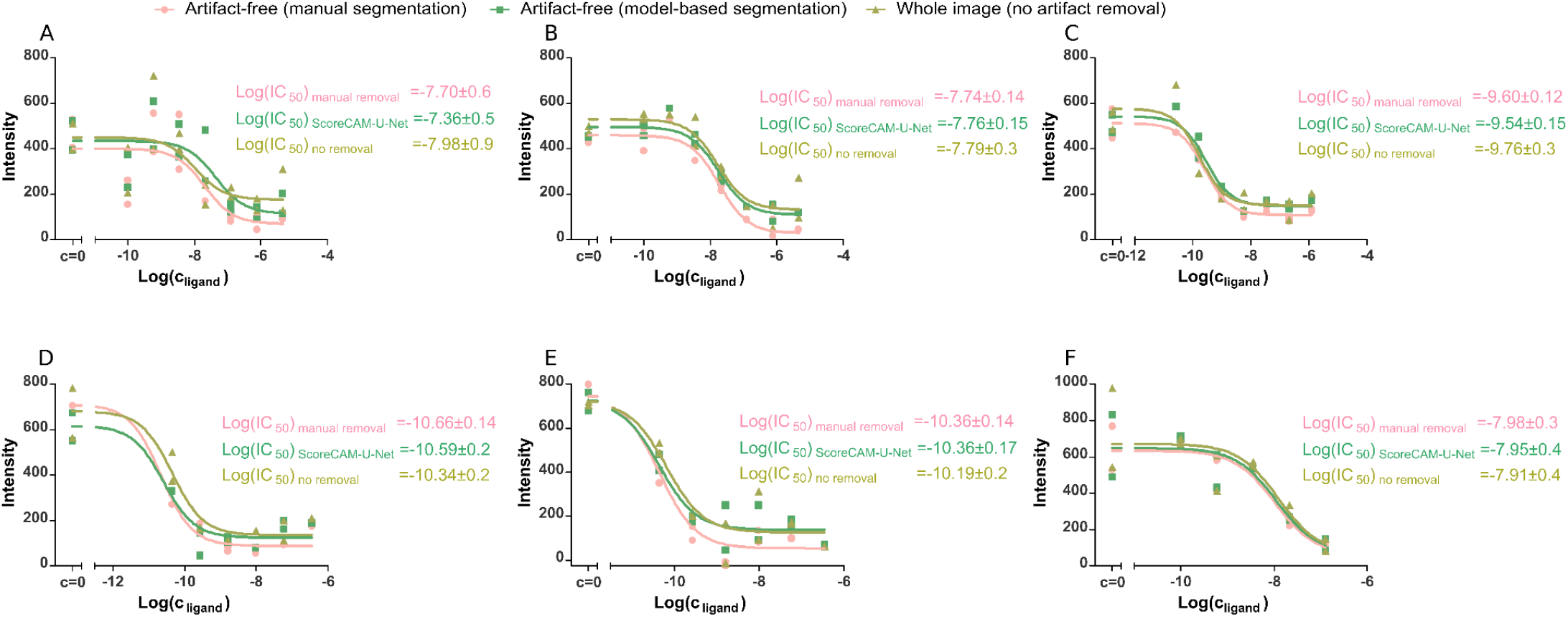
Cell fluorescence intensity dependence on M4 receptor ligand concentration determined with live-cell fluorescence microscopy at the presence of 2 nM UR-CG072. Displacement curves of three different ligands are shown: UNSW-MK259 (**A, B** and **F**), atropine (**D** and **E**) and UR-SK75 **(C)**. Three different artifact removal methods at the image analysis stage are compared (colors): manual artifact segmentation, ScoreCAM-U-Net segmentation and no artifact removal. For each combination of ligand and artifact removal method a regression analysis is performed with Hill equation (Hill coefficient fixed at 1) with the best fits shown as continuous lines. Each displacement curve was measured in duplicates with each data point representing the average fluorescence intensity of cells in each well.

